# Simulating multi-level dynamics of antimicrobial resistance in a membrane computing model

**DOI:** 10.1101/306100

**Authors:** Marcelino Campos, Rafael Capilla, Fernando Naya, Ricardo Futami, Teresa Coque, Andrés Moya, Val Fernandez-Lanza, Rafael Cantón, José M. Sempere, Carlos Llorens, Fernando Baquero

**Author notes:** To whom correspondence should be addressed: Fernando Baquero, Department of Microbiology, Ramón y Cajal University Hospital, Carretera de Colmenar km 9,1.28034 Madrid, Spain., and Carlos Llorens, 2Biotechvana, Valencia, CEEI Building, Benjamin Franklin Av. 12, Valencia Technological Park, 46980 Paterna, Spain.

## Abstract

Membrane Computing is a bio-inspired computing paradigm, whose devices are the so-called membrane systems or P systems. The P system designed in this work reproduces complex biological landscapes in the computer world. It uses nested “membrane-surrounded entities” able to divide, propagate and die, be transferred into other membranes, exchange informative material according to flexible rules, mutate and being selected by external agents. This allows the exploration of hierarchical interactive dynamics resulting from the probabilistic interaction of genes (phenotypes), clones, species, hosts, environments, and antibiotic challenges. Our model facilitates analysis of several aspects of the rules that govern the multi-level evolutionary biology of antibiotic resistance. We examine a number of selected landscapes where we predict the effects of different rates of patient flow from hospital to the community and *viceversa*, cross-transmission rates between patients with bacterial propagules of different sizes, the proportion of patients treated with antibiotics, antibiotics and dosing in opening spaces in the microbiota where resistant phenotypes multiply. We can also evaluate the selective strength of some drugs and the influence of the time-0 resistance composition of the species and bacterial clones in the evolution of resistance phenotypes. In summary, we provide case studies analyzing the hierarchical dynamics of antibiotic resistance using a novel computing model with reciprocity within and between levels of biological organization, a type of approach that may be expanded in the multi-level analysis of complex microbial landscapes.

## Introduction

Antibiotic resistance is the result of the complex interaction of discrete evolutionary entities placed in different hierarchical levels of biological organization, including resistance genes, mobile genetic elements, clones, species, genetic exchange communities, microbiomes, and hosts of these bacterial ensembles placed in particular biological environments (1,2,3). Under the influence of external environmental variation (such as exposure to antibiotics) each one of these evolutionary entities might have independent rates of variation and selection, but as they are hierarchically-linked, the changes in each one of them can influence all other entities (4), as they constitute a global “nested biological system” (5).

Membrane-computing is an individual-based natural computing paradigm aiming to abstract computing ideas and models from the structure and the functioning of living cells, as well as from the way the cells are organized in tissues or higher order structures (6,7). A kind of computational models using this paradigm are “P systems”, consisting in placing objects (in our case biological entities) into virtual cell-like or tissue-like membrane structures, so that one membrane or one cell (respectively) represents a hierarchical level, a region of the embedded system. For instance, each bacterial cell is a membrane containing plasmids (as objects), and a plasmid is a membrane containing genes (as objects). The mobility of entities, objects, across membranes is possible according to pre-established rewriting rules, and the collection of multisets of entities will evolve in a synchronous, parallel, and non-deterministic manner. The objects have assigned rules to pass through membranes (to mimic intracellular or intercellular transmission (8,9), to dissolve (to mimic elimination), and to divide themselves (to mimic replication). In this work, we use a P system to simulate multi-level dynamics of antibiotic resistance, based on our first published prototype (8,9). This computational model facilitates an approach that is computationally hard to accomplish or simply impossible to address experimentally. Our work allows the estimation and evaluation of global and specific effects on the frequency of each one of the biological entities involved in antibiotic resistance occurring because of changes taking place (as following antibiotic exposure) in one or (simultaneously) in several of them. Note that albeit antibiotic resistance is a major problem in Public Health, in terms of biosystems it is only a particular example of “evolution in action”. Our model can be easily applied to many other complex evolutionary landscapes, involving other genes, phenotypes, cells, populations, communities and ecosystems.

## Results

The main objective of the present work is to present the possibilities of membrane computational modeling as a powerful tool in the evaluation of the factors that, at various biological levels, might influence the dynamics of antibiotic resistance. The results provided below should not be taken as predictions of the evolution of resistance, just as illustrations of some of the possibilities of this model to study the multi-level dynamics of resistance, by simultaneously changing parameters in state variables and observing after a single run the effect in the frequency of resistant species and populations. Note that the model is probabilistic and the rules are selected in a probabilistic way. So, each computation produces an output in such manner that the results obtained are not entirely identical in consecutive runs of the program, but they are relatively close (see Fig SI1). In the next paragraphs, antibiotics (Ab) and the corresponding resistances (R) are named AbA, AbC and AbF, and AbAR, AbCR, and AbFR respectively; to facilitate reading, we suggest the identification of AbA as the Aminopenicillins, AbC as Cefotaxime-Ceftazidime, and AbF as Fluoroquinolones, using the initials of three of the major groups of antibiotics used in clinical practice (Table 1).

### The basic scenario in the hospital and community compartments

#### Dynamics of bacterial resistance phenotypes in *E. coli*

Waves of successive replacements of resistance phenotypes in hospital-based *E. coli* during 20,000 time-steps (about 2.3 years, as the time-steps represent approximately 1 hour/step) are illustrated in Fig 1. The main features of this process, mimicking clonal interference, are: 1) sharp decrease in the density of the fully susceptible phenotype (pink line); 2) rapid increase of the phenotype AbAR, aminopenicillin resistance, resulting from the transfer of the plasmid with AbAR to the susceptible population, and consequent selection (red); 3) increase by selection, and marginally by acquisition of mutational resistance, of the phenotype AbFR, fluoroquinolone resistance (violet); 4) increase of double resistances AbAR and AbFR, by acquisition of an AbFR mutation with the organisms of AbAR-only phenotype, and by the transfer of the plasmid encoding AbAR from the AbAR-only phenotype to the AbFR-only phenotype (brown); 5) increase of the phenotype with double resistances AbAR and AbCR by capture by the AbAR-only predominant phenotype of a plasmid containing AbCR, cefotaxime resistance that originated in *K. pneumoniae* (light blue); 6) almost simultaneous emergence but later predominance of the multi-resistant organisms with phenotype AbAR, AbCR, and AbFR by mutational acquisition of AbFR by the double-resistant phenotype AbAR-AbCR and, also, of the plasmid-mediated AbCR by the AbAR-AbFR phenotype (dark blue); 7) close in time, emergence, but with low density, of the phenotype AbCR-only, by the acquisition of the plasmid encoding AbCR by the fully-susceptible phenotype and the AbAR phenotype, and loss of plasmid-mediated AbAR by incompatibility with the incoming plasmid (light green); 8) the acquisition of the AbFR mutation by the AbCR-only phenotype, or by plasmid-reception of an AbCR trait from *K. pneumoniae* in AbFR, giving rise to the phenotype AbCR-AbFR (olive green). In the community, where the antibiotic exposure is less frequent, a similar dynamic sequence occurs, but at a much slower rate (fig. 2).

**Figure 1.**
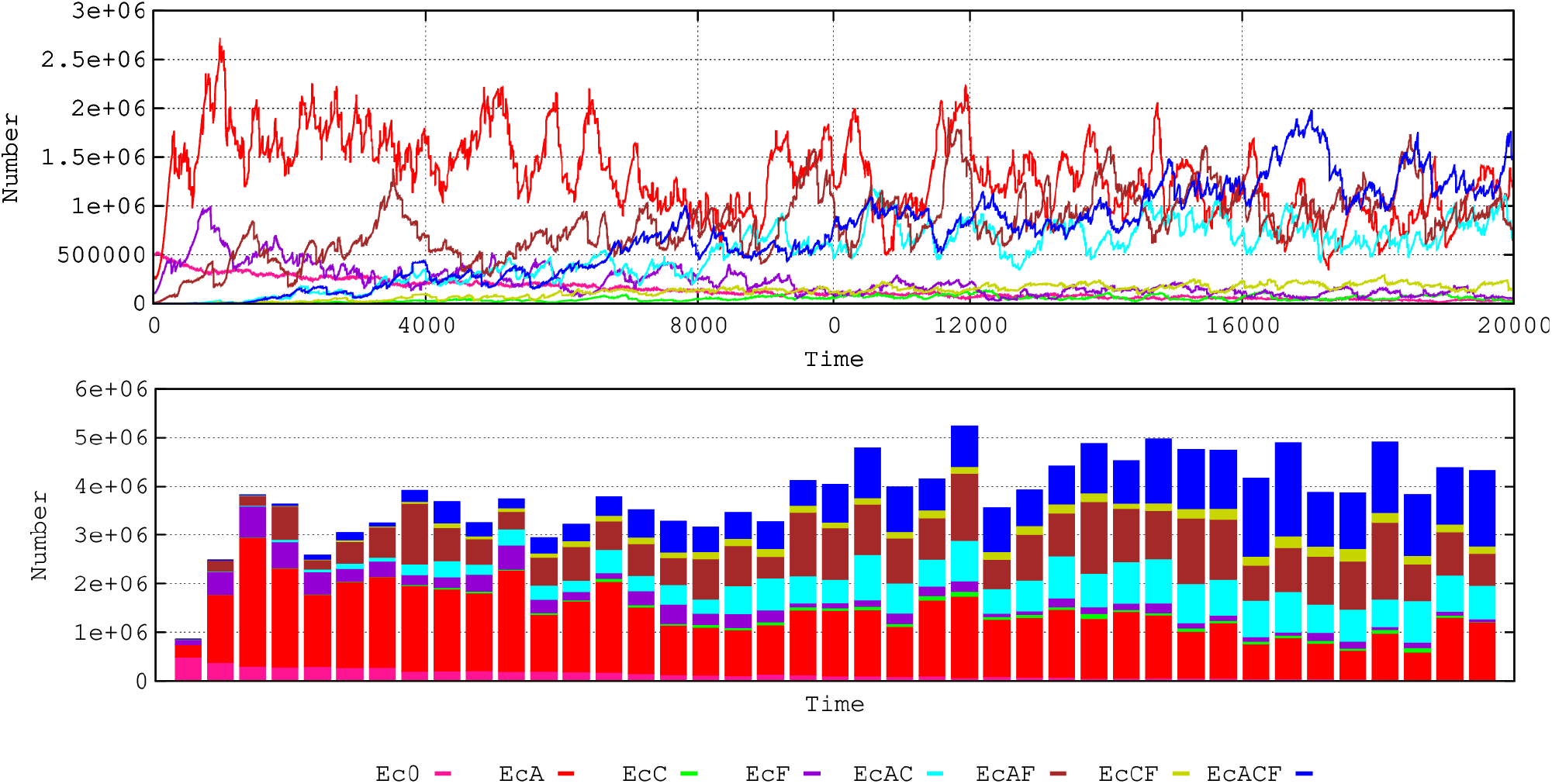
Dynamics of bacterial resistance phenotypes in *E. coli*. Pink, susceptible; red, AbAR (AMP); violet, AbFR (FLQ); brown, AbAR and AbFR; light blue, AbAR and AbCR; dark blue, AbAR, AbCR and AbFR; light green, AbCR; olive green, AbCR and AbFR. In ordinates, number of hecto-cells (h-cells, packages of 100 identical cells) in all hosts-ml (each host represented by 1 ml of colonic content); in abscissa, time (1000 steps, roughly equivalent to 42 days).

#### Dynamics of bacterial species

Antibiotic use and antibiotic resistance influence the long-term dynamics of bacterial species in hospital environment (Fig. 2 C, D). In the conditions of our basic scenario, *E. coli* populations (black) tend to prevail. *E. faecium* (violet) and *K. pneumoniae* (yellow-green) populations were maintained along the experiment. In the community, *E. coli* has a stronger dominance over other species, and similar dynamics occur as in the hospital, at slower rates.

**Figure 2.**
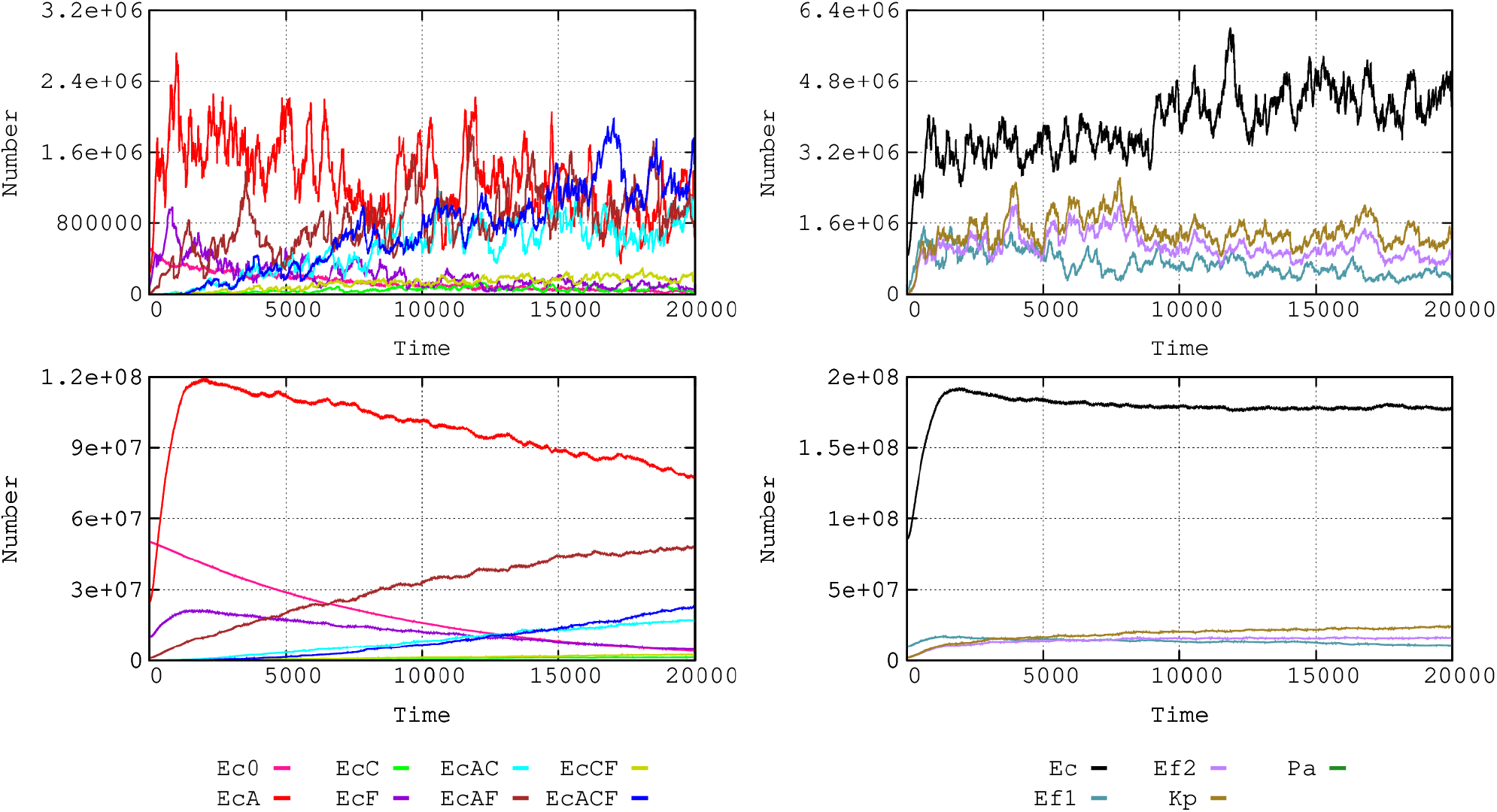
Comparative dynamics of *E. coli* phenotypes in the hospital (A), and the community (B); axes and color code, as in Fig 1. Species dynamics in the hospital (C) and the community (D): *E. coli* (black), *K. pneumoniae* (yellow green), *E. faecium* AbAS (violet), and *E. faecium* AbAR (dark green). *P. aeruginosa* is not visible in this representation (low numbers).

*Klebsiella pneumoniae* (Fig SM3) is intrinsically resistant to AbA and, in our case, it harbors a plasmid encoding AbCR (CTX), and a mutation encoding AbFR (FLQ). In the hospital, the AbCR phenotype is readily selected. However, because of the high density of *E. coli* with the plasmid-mediated AbAR, several *Klebsiella* strains receive this plasmid. These *Klebsiella* strains have no benefit from this plasmid because they are intrinsically aminopenicillin-resistant, but incompatibility with the plasmid determining AbCR occurs, eliminating AbCR from the recipients and giving rise to the phenotype AbAR-AbFR (purple). That contributes to the decline in AbCR-containing phenotypes (olive green). In any case, the dominance of *E. coli* prevents a significant growth of *K. pneumoniae. Enterococcus faecium* (Fig SM3) is intrinsically resistant to AbC (AbCR, CTX), but there are two variants, one AbA (AMP) susceptible, and the other resistant, this last one has also AbFR. However, the AbAS variant can acquire the AbAR trait from the resistant one by (infrequent) horizontal genetic transfer and becomes an AbAR donor. There is replacement dynamics of AbAS by the AbAR phenotype.

#### Influence of baseline resistance composition on the dynamics of bacterial species

The local evolution of antibiotic resistance can depend on the baseline composition of susceptible and resistant bacterial populations (Fig 3). In a baseline scenario, we consider a density of 8,600 h-cells (1 h-cell=100 identical cells, see the section “quantitative structure of the basic model application” below) of *E. coli* of which 5,000 are susceptible, 2,500 have plasmid-mediated aminopenicillin-resistance (PL1-AbAR), 1,000 have fluoroquinolone resistance (AbFR), and 100 combines both resistances. To mimic a “more susceptible scenario,” values were changed to 8,000 susceptible, 500 with PL1-AbAR, 50 with AbFR, and 50 with PL1-AbAR and AbFR. A higher proportion of susceptible *E. coli* facilitates the increase of the more resistant organisms, *K. pneumoniae* and AbAR *E. faecium*. Because of the selection of *K. pneumoniae* (olive green) harboring cefotaxime-resistance (PL1-AbCR), and the ability of transfer of the PL1 plasmid to *E. coli*, the proportion of *E. coli* with cefotaxime-resistance (mainly light and dark blue) increases in the scenario with a lower resistance baseline for *E. coli*. This example illustrates the hypothesis that a higher prevalence of resistance in the *E. coli* component of the gut flora might reduce the frequency of other resistant organisms, which might inspire interventions directed to restore susceptibility in particular species (10, 11).

**Figure 3.**
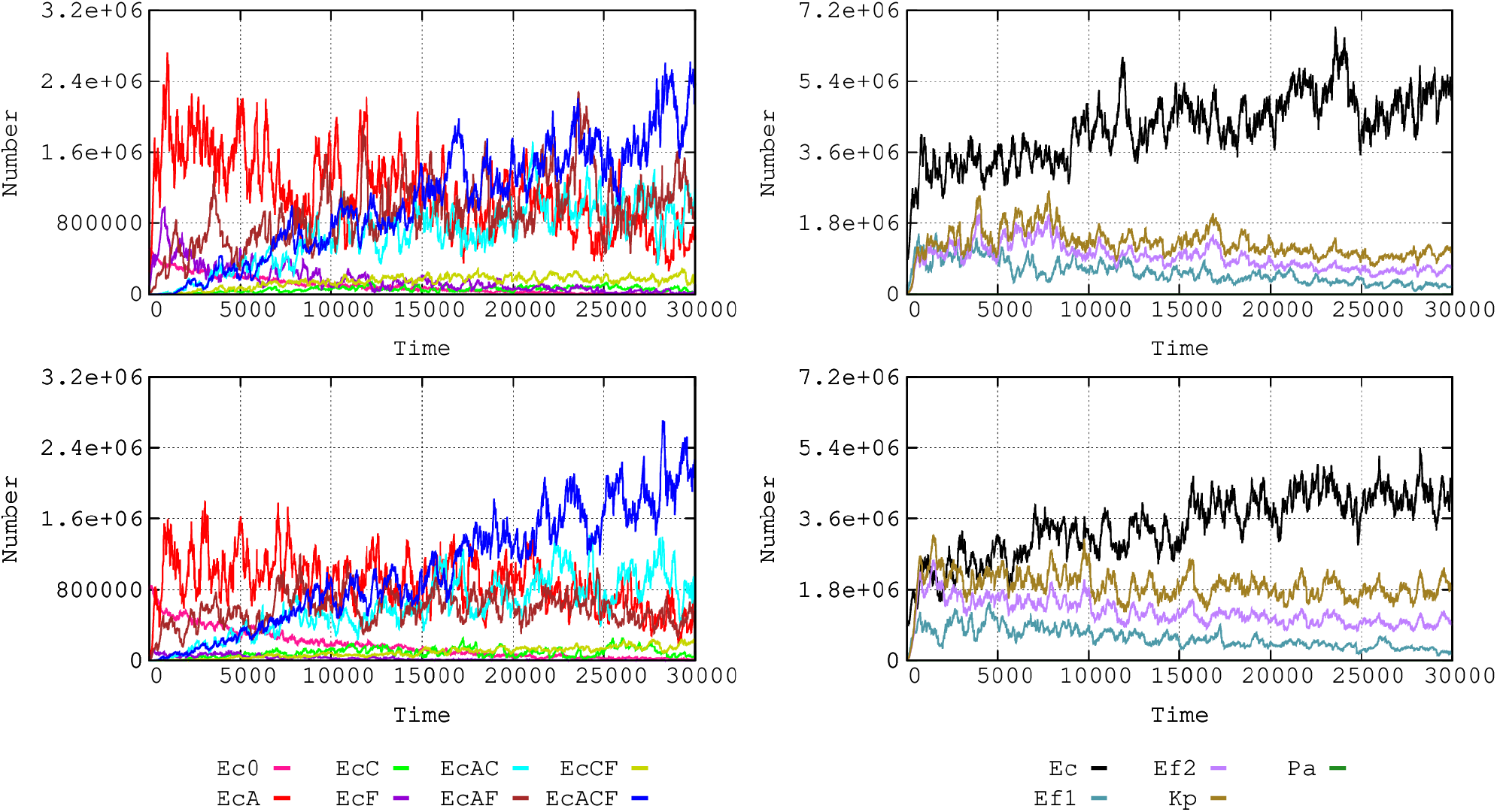
Influence of baseline *E. coli* resistance phenotypes composition on the dynamics of bacterial species. On the left, comparative dynamics of *E. coli* phenotypes in the basic hospital scenario (up) and with reduced numbers of resistant phenotypes (down). Colors and axes, as in Fig. 1. On the right, comparative dynamics of bacterial species in the basic model (up), and the reduced basal resistances (down); colors as in Fig 2.

#### Single clone *E. coli* dynamics: influence of baseline resistances

In the previous analysis, subpopulations of *E. coli* were characterized by their antibiotic-resistance phenotype (phenotype populations). Alternatively, we can follow the evolution of four independent *E. coli* clones, each one tagged in the model with particular signals (unrelated with AbR), Ecc0, EccA, EccF, EccAF (see Table 1), and starting with specific resistance traits, allowing for the possibility that the frequency of these “ancestor clones” may change through time within a clone by the gain or loss of a trait. Figure 4 shows the densities of these ancestor clones along time. The detail of sequential trait acquisition for each one of these clones is shown in Fig SM2. The fully susceptible *E. coli* clone (Ecc0) first acquires AbAR (red), and AbCR (green). The AbAR phenotype facilitates the capture by lateral gene transfer of AbCR (CTX), giving rise to the double AbAR-AbCR phenotype (light blue). The incorporation of AbF-R (violet, FLQ) in the fully susceptible clone occurs early, later in the AbAR population, so that the rise of the multi-resistant phenotype (dark blue) occurs later and again at low numbers. The presence of the AbAR trait in the clone at time 0 (EccA) increases the success of the clone, including the acquisition of AbFR, and the multi-resistant phenotype. Interestingly, the presence of AbFR (fluoroquinolones-R) at the origin (EccF), was critical to enhance the numbers of double-resistant and multi-resistant phenotypes. The clones that were more susceptible at the origin remain relatively stable in numbers, suggesting that clonal composition tends to level-off along the continued challenges under antibiotic exposure.

**Figure 4.**
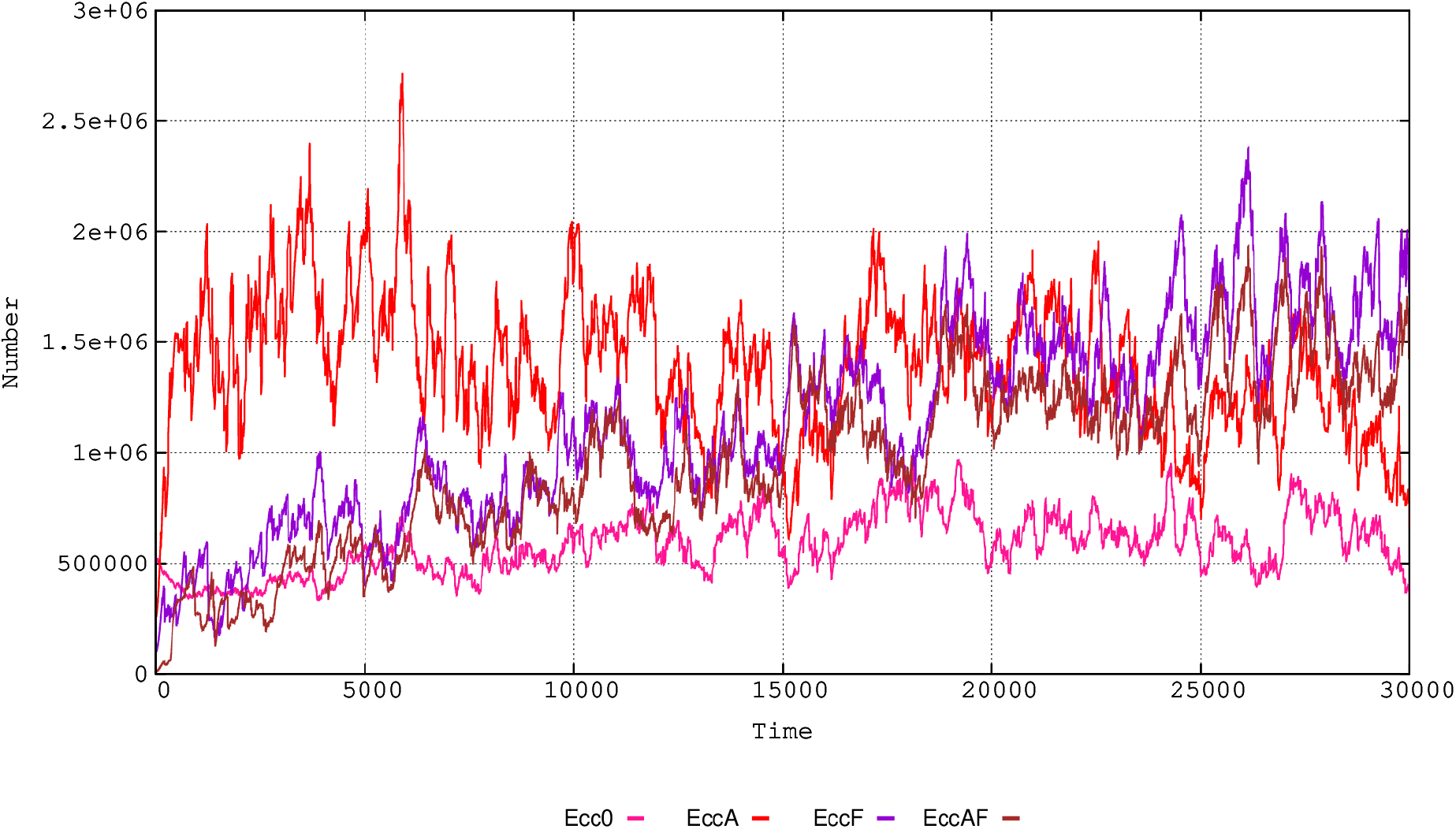
Single clone *E. coli* dynamics in the hospital: influence of baseline resistances. In pink, clone Ecc0 starting with full susceptibility, in red, with AbAR (EccA); in violet, with AbFR (EccF)); in brown, with AbAR and AbFR (EccAF).

#### Dynamics of mobile genetic elements and resistance traits

We consider *E. coli, K. pneumoniae*, and *P. aeruginosa* as members of a “genetic exchange community” (12,13) for the plasmid PL1. In Fig. 5, we can compare the evolutionary advantage of the same resistance phenotypic trait (AbAR) when harbored in a plasmid, as in *E. coli* or in the chromosome, as in *K. pneumoniae*. The overall success of the plasmid PL1 (blue line) benefits from the fact that this mobile element is selected by two different antibiotics (AbA and AbC, resistance shown in red and green lines respectively). Interestingly, resistance to AbFR (violet) is selected from early stages of the experiment, and after 4,000 steps it converges with the AbCR, a plasmid-mediated trait, meaning that this plasmid is maintained almost exclusively in strains harboring AbFR gene, similar to empirical findings (14,15). If the conjugation rate of PL1 was increased, the main effect would be the reduction in selection of *K. pneumoniae*, as the predominance of the PL1-AbAR plasmid from the more abundant populations of *E. coli* tended to dislodge PL1-AbCR from *K. pneumoniae* (results not shown).

**Figure 5.**
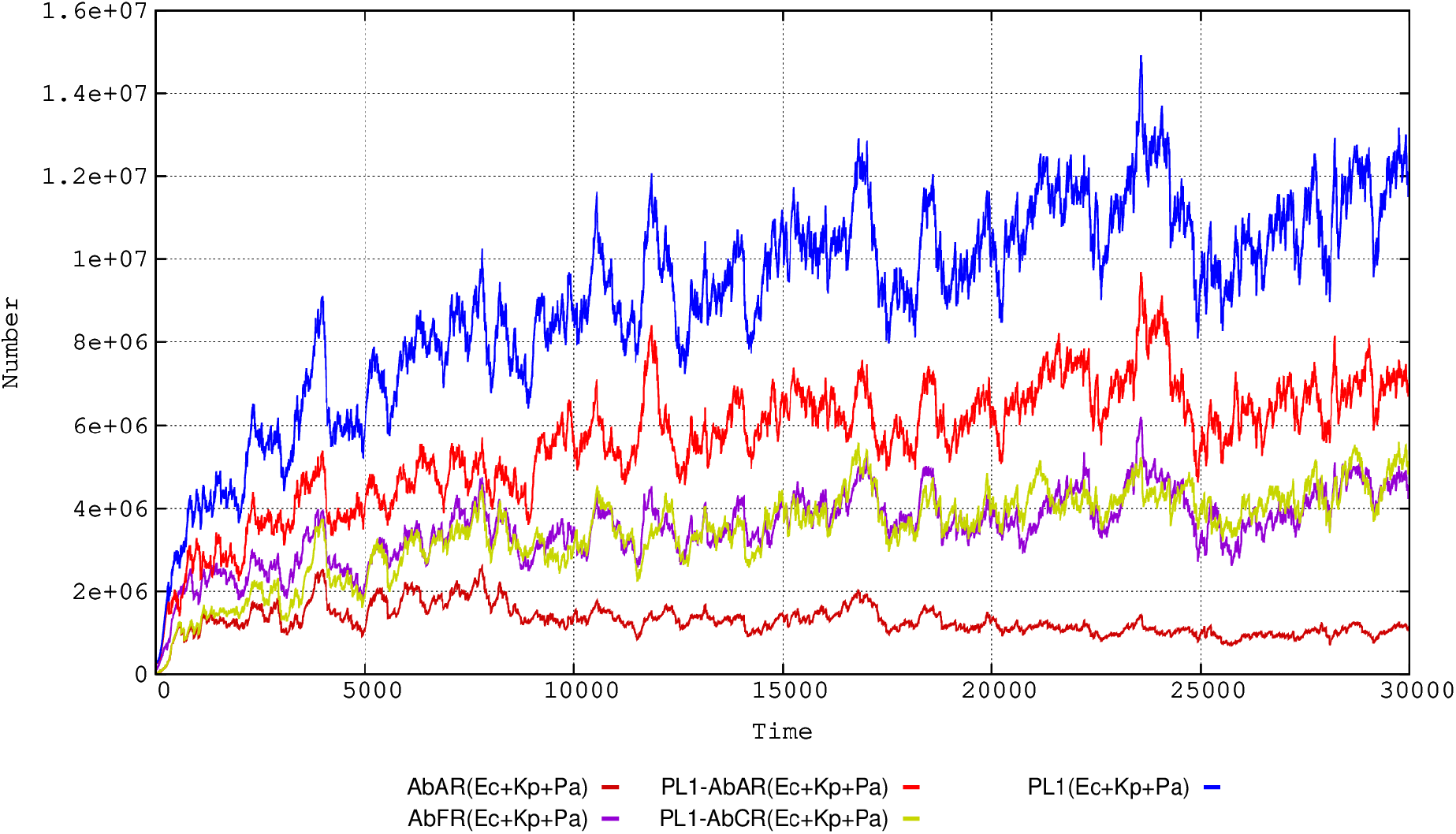
Dynamics of a plasmid and resistance traits in the hospital environment. The species *E. coli, K. pneumoniae* and *P. aeruginosa* are included as a genetic exchange community. In blue, total number of the plasmid PL1; in bright red, plasmid PL1 with the gene AbAR (AMP); in green, PL1 with AbCR (CTX); in violet, chromosomal AbFR (FLQ) gene; in red-brown, chromosomal AbAR (as in *K. pneumoniae*). In ordinates, number of plasmids or resistance traits in h-cells (packages of 100 identical cells) in all hosts-ml (each host represented by 1 ml of colonic content).

### Dynamics under changing scenarios in the hospital and community compartments

#### Frequency of patient flow between hospital and community

The frequency of exchange of individuals between the hospital and the community (hospital admission and discharge rates) influences the evolution of antibiotic resistance (Fig 6). This occurs because sensitive bacteria enter the hospital with newly admitted patients from the community (where resistance rates are low), and this “immigration” allows sensitive bacteria to “wash out” resistant bacteria (16). Multi-resistant *E. coli* strains emerge much earlier with decreased flow rates, as bacteria resistant to individual drugs have more time to coexist and thus exchange resistances by gene flow, and because the length of “frequent exposure” to different antibiotics increases, and consequently selection (17). The effect of slow flow of patients to the community is a late reduction in multiresistance (AbAR-AbCR-AbFR) and earlier double resistances (AbAR-AbFR) and (AbAR-AbCR). In the community compartment, however, multi-resistance increases when the flow from the hospital is more frequent (4 h).

**Figure 6.**
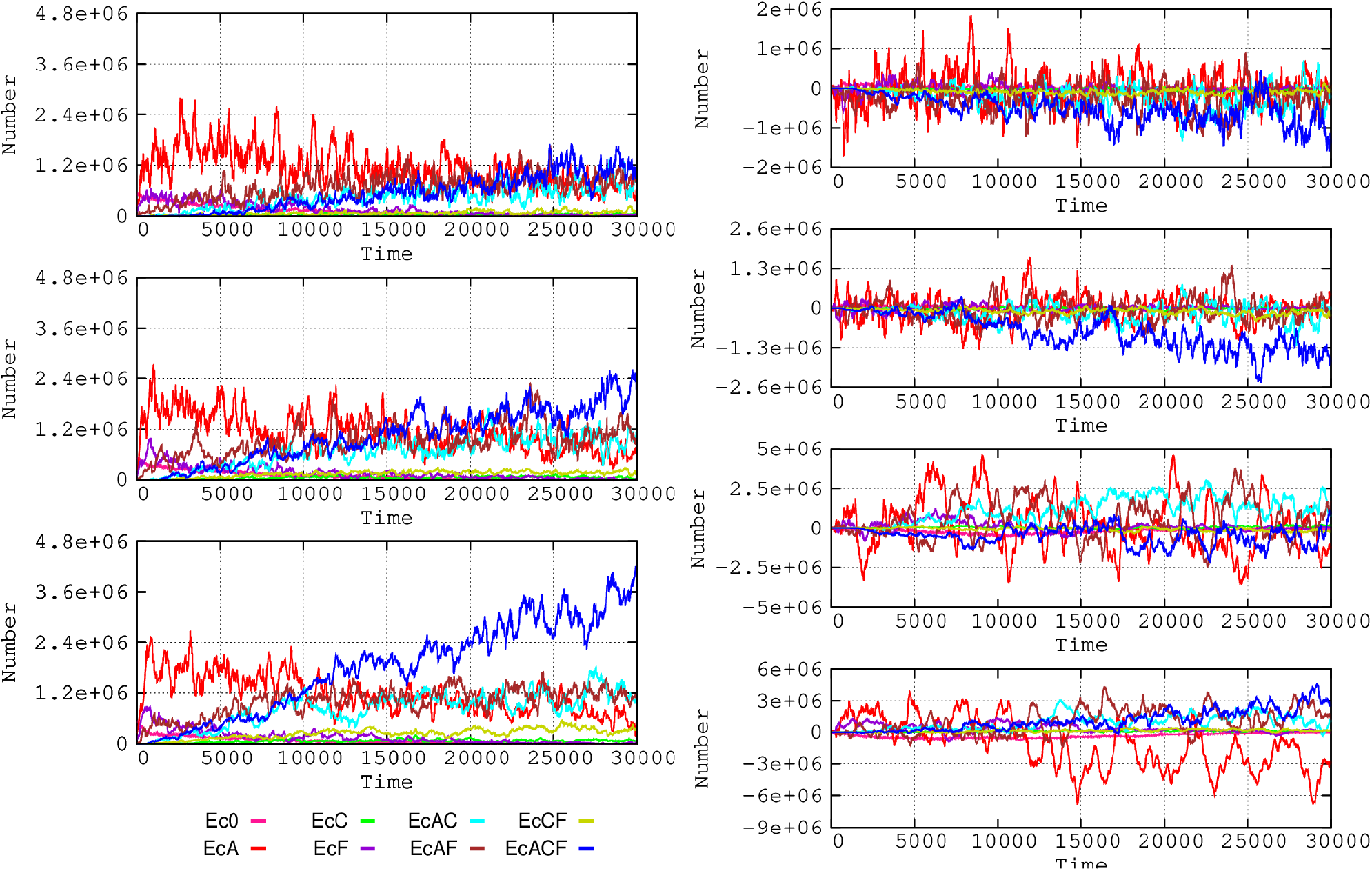
Influence of patients’ flow between hospital and community. On the left, influence on *E. coli* resistance phenotypes in the hospital when one patient is admitted at/discharged from the hospital every 2 (top), 4 (middle), or 8 hours (bottom).

#### Frequency of patients treated with antibiotics

The proportion of patients exposed to antibiotics increases selection of antibiotic resistance (16). We analyzed this effect in our model considering proportions of 20%-10%-5% of patients exposed to 7 consecutive days of antibiotic therapy, three doses per day (Fig 7). If a high proportion (20%) of patients are treated, *E. coli* multi-resistance is efficiently selected, as well as resistant *K. pneumoniae* and *E. faecium*. If this proportion is reduced to 10%, and particularly to 5%, there is a strong reduction in the amount of resistant *E. coli* cells and the emergence of multi-resistant bacteria is delayed (individual resistance data not shown for these species). However, the evolution of *E. coli* towards more multiresistance partially counteracts the selective advantage of these species, restricting their growth to some extent, even in the scenario of high density of treated patients.

**Figure 7.**
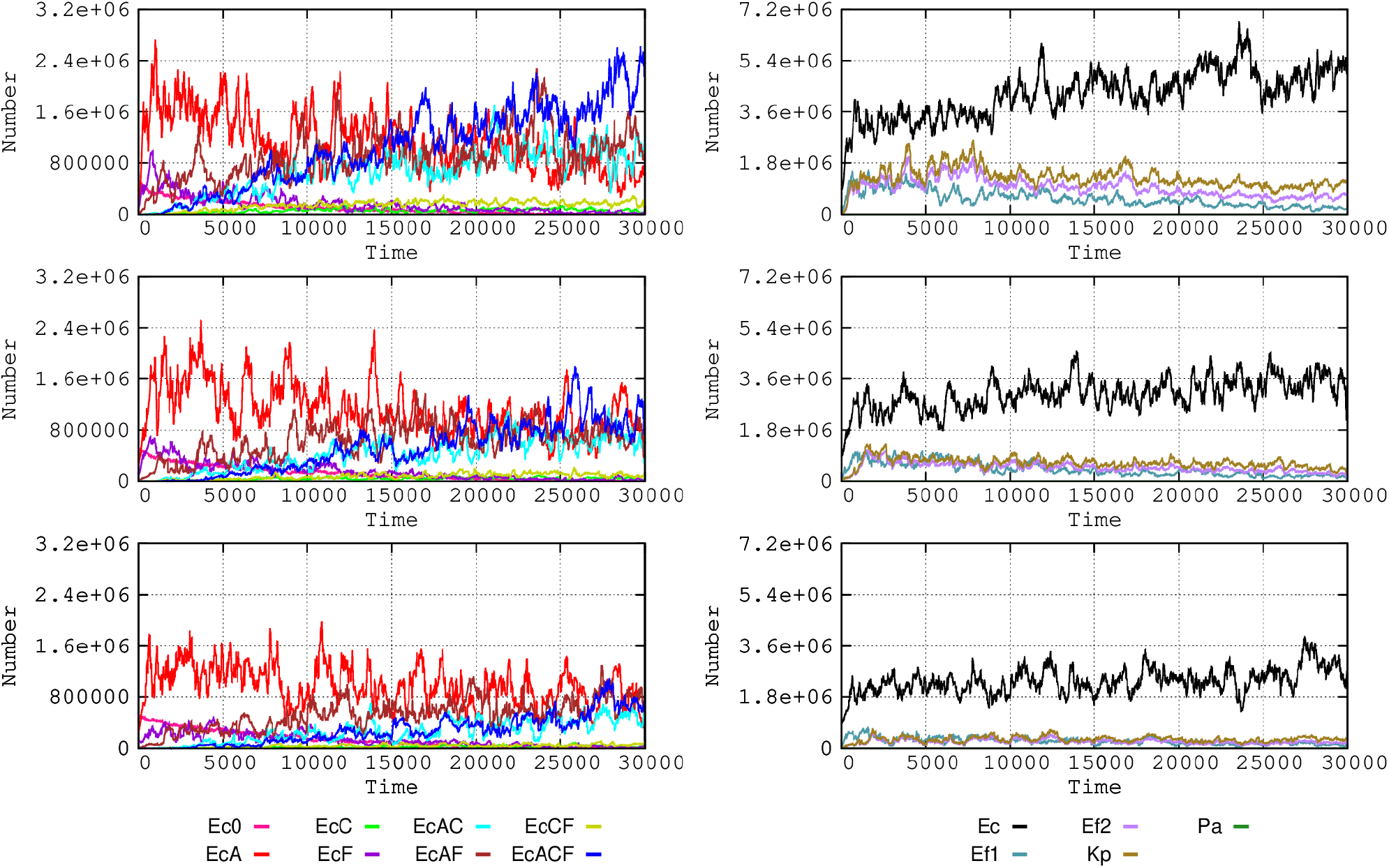
Influence of the frequency of patients treated with antibiotics. On the left, *E. coli* phenotypes when 20% (up), 10% (mid) or 5% (down) of patients receive antibiotics during a week, three doses per day. In the right part, effect on bacterial species. Colors as in Figs 1 and 2.

#### Frequency of bacterial transmission rates in the Hospital

Transmission of bacteria (any type of bacteria, including commensals) among individuals in the hospital influences the spread of antibiotic resistance. The effect of transmission rates of 5%, and 20% per hour was analyzed (Figure 8), expressing the proportion of individuals that acquire any kind of bacteria from another individual per hour. These rates might appear exceedingly high, indicating very frequent transmission between hosts, but we refer here to cross-colonization rate involving “any type of bacteria”. Normal microbiota transmission rates between hosts have never been measured, probably requiring a complex metagenomic approach (18). Differences in evolution of *E. coli* phenotypes comparing 10% and 20% of colonization rates are unclear; maybe 10% transmission produce full effects, and 20% does not add much more. The subtractive representation allows discernment of a global advantage for the multi-resistant phenotypes (AbAR-AbCR-AbFR) when the proportion of inter-host transmission rises from 5 to 20%. The mono-resistant AbAR phenotype tends to be maintained longer under low contagion rates. Note that multi-resistant phenotype “bursts” occur (dark blue spikes in the figure) also with low contagion rates (5% box in fig. 8), and “bursts” of less-resistant bacteria (red spikes) also occur in high contagion rates (20% box). It is to be noticed that the increase in cross-colonization rates favors not only the transmission of resistant populations, but also of the more susceptible ones, in a certain extent compensating the spread of the resistant phenotypes populations.

**Figure 8.**
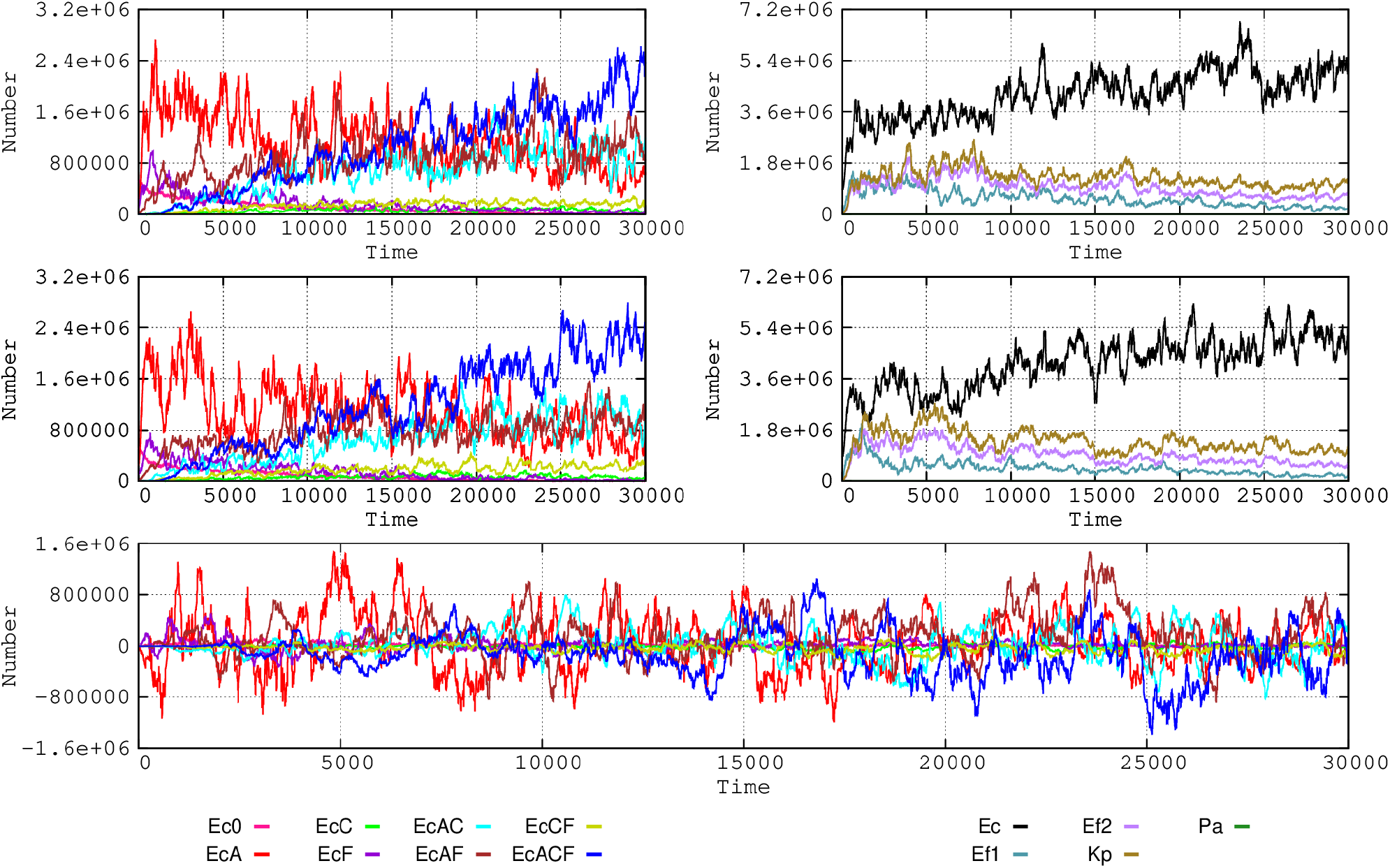
Influence of the frequency of bacterial cross-transmission rates in the hospital. On the left, dynamics of *E. coli* phenotypes when bacterial exchanges between patients occur in 5% (up) or 20% (down) per hour. A subtractive representation is provided below (5 vs. 20%). On the right, influence on the species composition: 5% (up), and 20% (down). Colors as in Figs 1 and 2.

#### Size of transmitted bacterial load

The absolute number of intestinal bacteria that are transmitted from one host to another one is certainly a factor influencing the acquisition of resistant (or susceptible) bacteria by the recipient. However, this number is extremely difficult to determine, as it depends not only on the mechanism of transmission (19,20), but also because the recipient might harbor bacterial organisms indistinguishable from those that are transmitted (21). On the other hand, efficient transmission able to influence colonic microbiota depends on the number of bacteria in the donor host, and the colonizing ability of different bacteria, not only in the intestine, but also in intermediate locations in the body, as probably the mouth or upper intestine (22). To show the potential effect of different bacterial loads acting as inocula, we consider a final immigrant population reaching the colonic compartment equivalent to 0.1%, 0.5% and 1% of the donor microbiota. As in previous cases, the evolution of multi-resistance favors *E. coli* (Fig SI4). Multi-resistant *E. coli* emerges earlier and reaches higher counts in higher-count inocula, but less resistant strains are maintained because the higher-count inocula also contain more susceptible bacteria.

#### Intensity of the effect of antibiotics on bacterial populations

The question of the relation of the “potency” (intensity of antibacterial activity) of antibiotics in relation with the selection of resistance has been a matter of recent discussions (23, 24, 25, 26). To illustrate the point, we changed the bactericidal effect of the antibiotics used in the model. Clinical species were killed at rates of 30%-15% (reflecting population decrease) the first and second hour of exposure respectively, and these rates were decreased to 7.5-3.75%. Note that these modest killing rates intend to reflect the diminished effect of antibiotics in slow-growing clinical bacteria located in a complex colonic microbiome. The more susceptible *E. coli* phenotypes are maintained for longer when the killing intensity of antibiotics is lower; on the contrary, the multi-resistant phenotype emerges earlier and reaches higher numbers when the intensity of antibiotic action increases (Fig 9). Under high antibiotic intensity, there is also a (small) increase in the resistant *K. pneumoniae* and *E. faecium* phenotypes. This experiment shows that a high rate of elimination of the more susceptible bacteria favors the colonization by the more resistant ones.

**Figure 9.**
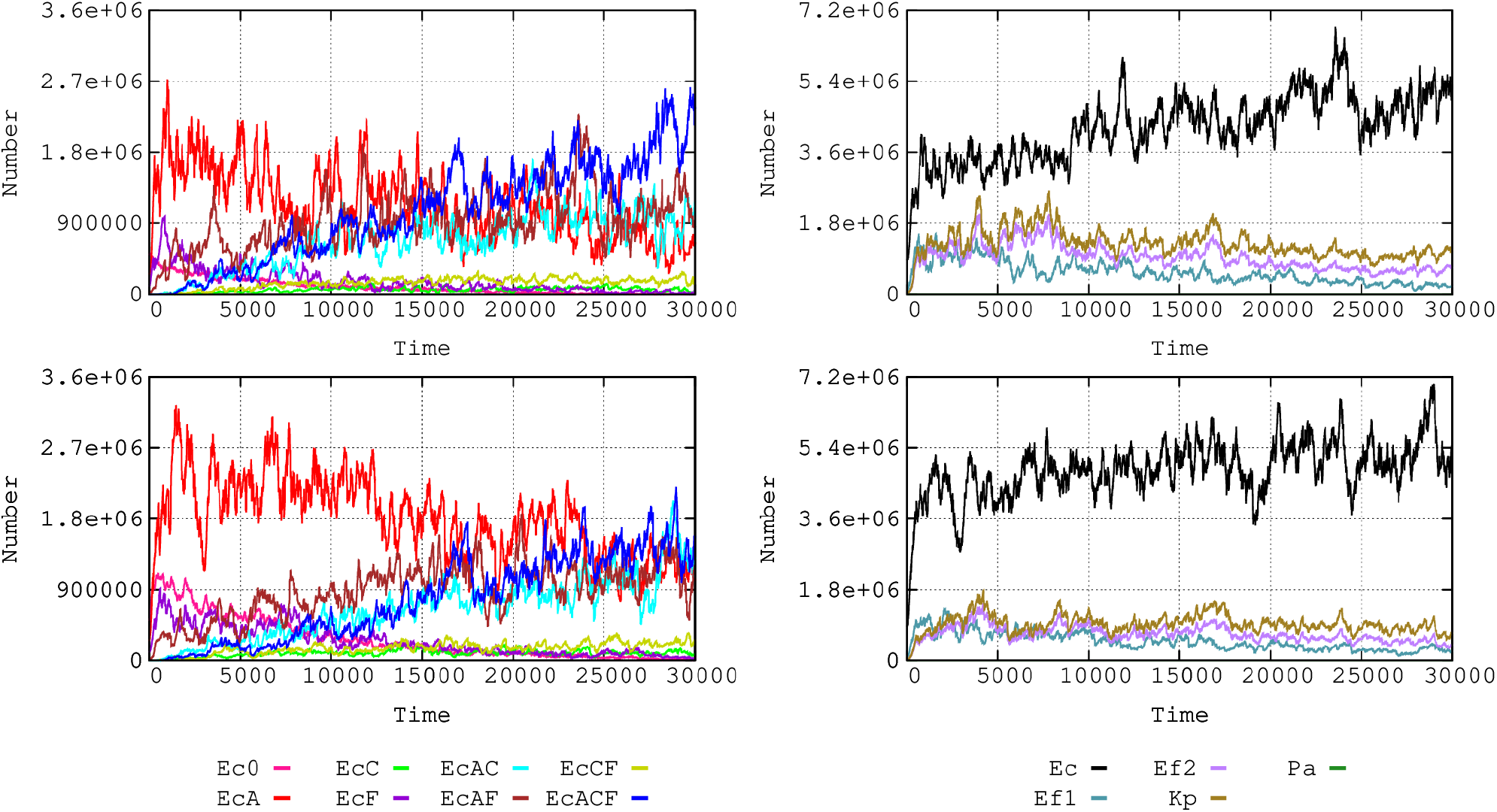
Influence of the activity of the antibiotic on *E. coli* phenotypes (left) and the species composition (right). Upper panels, susceptible bacteria are eliminated 30% after the first hour of exposure and 15% after the second hour; in the lower panels, the elimination is lower, 7.5% the first hour and 3.75% the second hour. Colors as in Figs 1 and 2.

#### Intensity of the antibiotic effect on colonic microbiota

The proportion of the colonic microbiota killed by antibiotic treatment, and thus the size of the open niche for other strains to multiply, constitutes an important factor in the multiplication of potentially pathogenic bacteria, and hence facilitates acquisition (mutational or plasmid-mediated) of resistance, and transmission to other hosts. In the basic model, reduction of the population is 25% for AbA, 20% for AbC, and 10% for AbF; in an alternative scenario these proportions were modified to 10%, 5% or 2% respectively. The result of this change is impressive (Figure 10): not only the number of bacteria is reduced but the evolution towards antibiotic resistance (EC) occurs at a slower rate, and even if the proportion of resistance phenotypes steadily increases along time, its absolute number does not grow, thus limiting host-to-host transmission.

**Figure 10.**
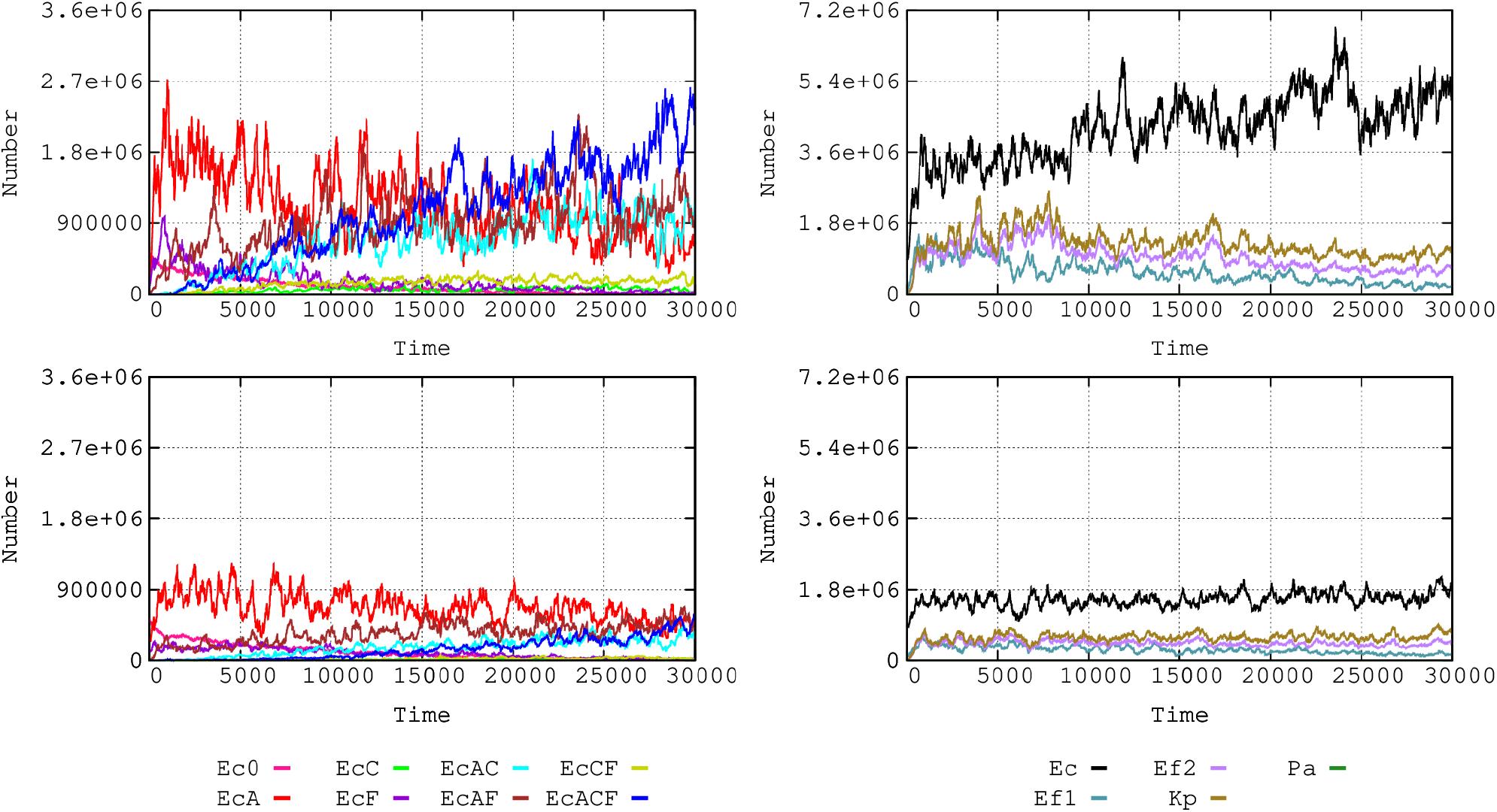
Influence of the intensity of the antibiotic effect on colonic microbiota of patients in the hospital. On the left, effects on *E. coli* phenotype of a reduction in microbiota of 25% for AbA, 20% for AbC, and 10% for AbF (upper panel); these values were reduced to 10%, 5% or 2% respectively (lower panel). The effects on the species composition is shown at the right side.

#### Strength of antibiotic selection on resistance traits

Strength of antibiotic selection is an important parameter in evolutionary biology of antibiotic resistance (27). Our computational model allows heuristic knowledge about the strength of selection of an antibiotic for a particular resistance trait, considering how the resulting trend is (or not) compatible with the observed reality. An example case is the unanswered question: -do plasmid-mediated cefotaxime-resistance (AbCR) also provides protection against aminopenicillins (AbAR)? Strains harboring TEM- or SHV-extended-spectrum beta-lactamases hydrolyzing cefotaxime probably retain sufficient levels of aminopenicillin hydrolysis to be selected by aminopenicillins. However, the phenotype cefotaxime-resistant/aminopenicillin-susceptible is rare in hospital isolates. In our model, this was investigated providing different strengths of ampicillin (AbA) selection for a cefotaxime-resistant phenotype (AbCR): no selection (0%), selection only in 10% of the cases (10%), and full selection (100%). The results of the model (Fig SM5) show that if ampicillin were able to select for cefotaxime-resistance the phenotype aminopenicillin-susceptible and cefotaxime-resistant should be prevalent from early stages. This is not what is observed in the natural hospital environment, suggesting that ampicillin is not a major selector for cefotaxime-resistance.

## Discussion

The rate of antibiotic resistance among bacterial species in a given environment is the result of the interaction of biological elements within a framework determined by many local variables, constituting a complex parameter space (28, 29, 30). There is a need to consider (in an integrated way) how changes in these parameters might influence the evolution of resistant organisms. This endeavor requires the application of new computational tools that should consider the nested structure of the microbial ecosystems, where mechanisms of resistance (genes) can circulate in mobile genetic elements among bacterial clones and species belonging to genetic exchange communities (12, 13) located in different compartments (as the hospital, or the community). A number of different factors critically influence the evolution of this complex system, such as antibiotic exposure (frequency of treated patients, drug dosages, the strength of antibiotic effects on commensal bacterial communities, the replication rate of the microbial organisms, as well as the fitness costs imposed by antibiotic resistance, the rate of exchange of colonized hosts between compartments with different levels of antibiotic exposure (hospital and community), or the rates of cross-transmission of bacterial organisms among these compartments. The challenge that we are addressing in this work is to simultaneously combine for the first time all these (and potentially more) factors in a single computing model to understand the selective and ecological processes leading to the selection and spread of antibiotic resistance. In comparison with available classic mathematical models that have been applied to the study of evolution of antibiotic resistance (31), the one we are discussing in this work is far more comprehensive in terms of the level of capture of the multi-level parametric complexity of the phenomenon. Note that results obtained with the model and presented here correspond only to a very limited number of possible “computational experiments”, chosen to show the possibilities of the model, but a virtually unlimited number of other experiments, with different combinations of parameters, are feasible *à la carte* with a user-friendly interface. In addition, our model can illustrate principles, generate hypothesis and guide and facilitate the interpretation of empirical studies (32, 33). Examples of these heuristic predictions are that resistance (less antibiotic effect) in colonic commensal flora can minimize colonization by resistant pathogens, the possible minor role of aminopenicillins in the selection of extended-spectrum beta-lactamases (AbCR), or the possibility of the presence of plasmids containing aminopenicillin-resistance in *K. pneumoniae*, phenotypically “invisible” as this organism has chromosomal resistance to the drug.

Our results are presented in terms of the ensemble of biological entities contained in the whole landscape (for instance the hospital), aggregated across individual hosts. This “pooling” approach, originated in ecological studies, has already been used in antibiotic resistance (34). Environments (as the hospital) are depicted as single “big world” units colonized by “big world populations”, including those with are antibiotic resistant but also the susceptible ones, which can limit the spread of resistance, in a sense “spreading health” (35). In this scenario, how might antibiotics modify the available colonization space? (36, 37). Our model includes the elimination of part of the global colonic microbiota with antibiotic use, favoring the colonization of resistant organisms, previously in minority.

In our computational experiments we can reproduce the successive “waves” of increasingly resistant phenotypes, mimicking the clonal interference phenomenon (38). We show that the speed and intensity of this process depends on the global resistance landscape and the density and phenotype of the bacterial subpopulations. Our model predicts that previous mutational ciprofloxacin-resistance facilitates fast evolution of multi-resistance by horizontal acquisition of resistance genes (14, 15). We also show that the long-term dissemination of chromosomally-encoded genes is by far less effective than the spread of traits encoded in transferable plasmids, even though some limitations are detectable because of plasmid incompatibility. A frequently overlooked aspect of antibiotic resistance suggested by our membrane computing experiments, is that probably the evolution of multi-resistance favors at long term some predominant species, as *E. coli*, where there is also an increasing benefit for the more resistant clones.

The consequences of changes in the transmission and treatment rates of the hospital and the community were also explored in our model. Several mathematical models have also investigated these changes (16, 37, 38, 39, 40, 41, 42, 43, 44, 45). Is clear that the effect of reducing patient discharges and admissions in the hospital increases the local rates of antibiotic resistance, but in our model, the proportion of antibiotic treated patients in the hospital has the stronger effect, stressing the importance of a precision-prescribed antibiotic therapy (44). The role of increasing rates of hospital cross-colonization also influences the rise of resistance, but this effect seems lower than expected, probably because higher transmission rates also assures transmission of the more susceptible antibiotic populations, a kind of “washing out” process of resistance, as the one that occurs when the community-hospital flow increases (16). The model also predicts that the “amount” of bacteria transmitted between hosts favors the ascent of antibiotic resistance. We considered another frequently overlooked factor: the consequences of “intensity” (aggressiveness) of the antibiotic therapy, because of frequent dosage and particularly in terms of its ability to reduce the colonic microbiota, and therefore “colonization resistance” for resistant opportunistic pathogens (47).

Precise data are not always easy to obtain, and the type of mathematical or computational models should influence the results of predictions (48). However, because of the functional analogy of membrane computing with the biological world, we hypothesize that the trends revealed in our computational model reflect general processes in the evolutionary biology of antibiotic resistance. If the model were fed with objective data extracted from a real landscape (which will be possible with a user-friendly interface), it could provide a reasonable expectation of the potential evolutionary trends in this particular environment and could support the adoption of corrective interventions (49). Validation of this computational model is the next necessary step; to this goal, we are developing an “experimental epidemiology” model where the parameters could be altered and measured (50), and also planning prospective hospital-based observations.

Finally, we would like to stress that the type of membrane-computing model that was applied in this work can be easily escalated or adapted to a variety of applications in systems biology (51,52), and particularly to understand complex ecological systems with nested hierarchical structures and involving microorganisms (53).

## Material and Methods

### Software implementation and computing model

All computational simulations were performed using an updated version of ARES (Antibiotic Resistance Evolution Simulator), which is the software implementation of a P system for modeling of antibiotic resistance evolution (8). This P system model works with objects and membranes distributed in different regions organized in a tree-like structure, as the P system classic model, but now with more specific rules: the “object rules” can modify an object (evolution rules) or move the object out, in, or between membranes, the “membrane rules” can move membranes out, in, or between regions that contain them as “object rules” and can dissolve and duplicate membranes. When a membrane is dissolved all the membranes and objects inside disappears. For duplication we can define which objects will be duplicated and which ones will be distributed; the membranes are always distributed. The implementation of our P system uses a stochastic to apply the rules, the rules being ordered by priorities and each rule has a “probability” to be applied. Other computational objects can be introduced, either to tag particular membranes, or to interact with the embedded membranes, for instance mimicking antibiotics, according to a set of pre-established rules and specifications. We obtain an evolutionary scenario including several types of nested computing membranes emulating entities such as: i) resistance genes, located in the plasmid, other conjugative elements or in the chromosome; ii) plasmids and conjugative elements transferring genes between bacterial cells; iii) bacterial cells; iv) microbiotas where different bacterial species and subspecies (clones) can meet; v) hosts containing the microbiotic ensembles; vi) environment(s) where the hosts are contained. The current version of ARES (2.0) that can be freely downloaded at https://sourceforge.net/projects/ares-simulator/. ARES 2.0 runs in any computer (is a java application) albeit it is highly recommendable to install it in at least a 4× 6 Core Server and 128 GB of RAM. The original ARES web site at http://gydb.org/ares offers sections with information about the rules and parameters currently used by ARES.

### Anatomy of the model application

The current application of the model was structured accordingly with the following composition: 1) compartments containing individual hosts at particular densities, mimicking a hospital (H) and a community environment (C); flux of individuals between both compartments occurs at variable rates, mimicking admission or discharge from the hospital. 2) clinically relevant bacterial populations colonizing these hosts, from the species, Ec, *Escherichia coli*; Ef, *Enterococcus faecium*; Kp, *Klebsiella pneumoniae*), and Pa, *Pseudomonas aeruginosa*. These populations diversify from their initial phenotype by acquisition of mutations and/or mobile genetic elements, plasmids PL1 and for Ec, Kc, Pa circulating in these species, or, in Ef, conjugative elements (CO1). The cell can maintain two copies of the plasmid PL1 (containing resistance to AbA (PL1-AbAR) or AbC (PL1-AbCR) but not more, so that when a third plasmid PL1 enters the cell, one of the three is stochastically removed. AbCR produces some degree of resistance to AbA, and we consider this antibiotic also selects, in 10% of the cases, cells containing the plasmid PL1-AbCR. CO1 is an Ef “plasmid-like” mechanism of transfer of chromosomal gene AbAR (CO1-AbAR); a single copy of CO1-AbAR exist in the receiving host. Acquired resistance (not intrinsic) to AbA (AbAR) is mediated by the acquisition of PL1 (or CO1), resistance to AbC (AbCR), by acquisition of PL1 containing the AbCR resistance determinant, and resistance to AbF (AbFR) by mutation. Note that our representations, for example, when Ec0 (susceptible) receives PL1 with AbAR it becomes EcA, if PL1 with AbCR becomes Ec2C, and when Ec0, Ec1 or Ec2 mutate to AbFR become EcF, EcAF3, and EcCF. The acquisition of PL1 with AbAR by EcCF or PL1 with AbCR by EcAF produces the multi-resistant strain EcACF.

### Quantitative structure of the basic model application

**Hospitalized hosts in the population.** The number of hosts in the hospital and community environments reflects an optimal proportion of 10 hospital beds per 1,000 individuals in the community (https://data.oecd.org/healtheqt/hospital-beds.htm). In our model, the hospital compartment has 100 occupied beds, and corresponds to a population of 10,000 individuals in the community.

**The admission and discharge rates from hospital** are equivalent, 3-10 individuals/10,000 population/day (http://www.cdc.gov/nchs/data/nhds/%201general/). In the basic model, 6 individuals from the community are admitted to the hospital and 6 are discharged from the hospital to the community per day (approximately at 4 hour-intervals). Patients are stochastically admitted or discharged, meaning that about 75% of the patients stay in the hospital between 6 and 9 days.

**The bacterial colonization space** of the populations of the clinical species considered here (Table 1) and other basic colonic microbiota populations is defined as the volume occupied by these bacterial populations. In natural conditions, the sum of these populations was estimated in 10^8^ cells per ml of the colonic content. Clinical species constitute only 1% of the cells in each ml, and have a basal colonization space of 1% of each ml of colonic content, 0.01 ml. In the next section is explained how these spaces are considered for counting populations in the model.

The ensemble of other microbiota populations is considered in our basic study model as an ensemble surrounded by a single membrane. The colonic space occupied by these populations can change because of antibiotic exposure. Along a treatment course (7 days) the antibiotics AbA, AbC, and AbF reduce the intestinal microbiota 25%, 20% and 10% respectively. As an example, if we consider that 10% of the basic colonic populations were eliminated by antibiotic exposure, their now empty space (0.1 ml), will be occupied by antibiotic resistant clinical populations, and by the colonic populations that have survived the challenge. In the absence of antibiotic exposure, the colonic populations are restored in two months to their original population size. Clinical populations are comparatively faster in colonizing the empty space.

### Populations’ operative packages and counts

To facilitate the process of model running, we consider that 10^8^ cells in nature is equivalent to 10^6^ cells in the model. In other words, one “hecto-cell” (h-cell) in the model is an “operative package” of 100 cells in the real world. Because of the very high effective population sizes in bacteria, these 100 cells are considered as a uniform population of a single cell type. A certain increase in stochasticity might occur because of using h-cells; however, run replicates do not differ significantly (fig SM1). Also for computational efficiency, we considered that each patient (in hospital) or individual (in the community compartment) is represented in the model by 1 ml of its colonized colonic space (about 3,000 ml) and is referred as a “host-ml”. Consequently, in most of the figures we represent our results as “number of h-cells in all hosts-ml”.

### Quantitative distribution of clinical species and clones

In the basal scenario, the distribution of species in these 1,000,000 cells, contained in 1 ml, is the following: for EC, 860,000 cells, including 500,000 susceptible cells, 250,000 containing PL1-AbAR, 100,000 with the AbFR mutation, and 10,000 with both PL1-AbAR and AbFR mutation; for EF, 99,500 AbA susceptible and 20,000 AbAR. For KP, 20,000, with chromosomal AbAR, PL1-AbCR and AbFR; and PA, 500 containing PL1-AbCR. At time 0, this distribution is identical in hospitalized and community patients.

### Tagging starting clone populations in *E. coli*

To be able to follow the evolution of particular lineages inside *E. coli*, four ancestral clones (Ecc) were distinguished, differing in the original resistance phenotype, Ecc0 as a fully susceptible clone, EccA harboring PL1 determining AbAR, EccF harboring AbFR, and with EccAF with PL1-AbAR, and AbFR (Table 1). At time 0 each one of these clones is tagged with a distinctive “object” in the model which remains fixed to the membrane, multiplies with the membrane, and is never lost. Each one of the daughter membranes along the progeny can alter its phenotype by mutation or lateral gene acquisition, but the ancestral clone will remain detectable.

### Multiplication rates

We consider the basal multiplication rate (=1) the one corresponding to Ec0, where each bacterial cell gives rise to two daughter cells every hour. Comparatively, Ef=0.85, Kp=0.9, and Pa=0.15. The acquisition of a mutation, plasmid of a mobile element imposes an extra cost of 0.03. Therefore, Ec0=1, EcA=0.97 (because of the cost of PL1-AbAR), EcC=0.97 (cost of PL1-AbCR), EcF= 0.97 (cost of mutation); EcAF=0.94 (PL1-AbAR and AbFR), Ef(1)=0.85, Ef(2)=0.79 (CO1-AbAR and AbFR), Kp=0.84 (PL1-AbAR and AbFR), and Pa with PL1-AbCR =0.12 (PL1-AbCR). The number of cell replications will be limited by the available space (see above).

**Transfer of bacterial organisms from one host to another one** is expressed by the proportion of individuals that can stochastically produce an effective transfer of commensal or clinical, susceptible or resistant bacteria to another one (contagion index, CI). If contagion is 5%, or CI=5, that means that from 100 patients, 5 “donors” transmit bacteria to 5 others “recipients” per hour. In the case of the basic scenario, CI=5 in the hospital and CI=1 in the community (all results with CI=0.01 are available on request). In the basic scenario, donors contribute to the colonic microbiota of recipient individuals with 0.1, 0.5 and 1% of their own bacteria. This inoculum does not necessarily reflect the number of cells transferred, but also reflects endogenous multiplication after transfer, as proposed in other models (54). In any case, crosstransmission is responsible for most new acquisitions of pathogenic bacteria (55).

**Frequency of plasmid transfer between bacteria** occurs randomly and reciprocally at an equivalent high frequency among Ec and Kp; in the basic model, the rate is 0.0001, one effective transfer occurring in 1 of 10,000 potential recipient cells. Plasmid transfer occurs at a lower rate, of 0.000000001 in the interactions of Ec and Kp with Pa. Conjugative-elements) mediated transfer of resistance among Ef occurs at a frequency of 0.0001, but Ef are unable to receive or donate resistance genes to any of the other bacteria considered. In the case of Ec and Kp plasmids we consider plasmid limitation in the number of accepted plasmids, so that if a bacterial cell with two plasmids receives a third plasmid, there is a stochastic loss of one of the residents or the incoming plasmid, but all three cannot coexist in the same cell.

**Mutational resistance** is only considered in the present version of the model for resistance to AbF, fluoroquinolones. Organisms of the model-targeted populations mutate to AbF at the same rate, 1 mutant every 10^8^ bacterial cells per cell division.

### Antibiotic exposure

In the basic model, 5%-10%-20% of the individuals in the hospital compartment are under antibiotic exposure each day, each individual being exposed (treated) for 7 days. In the community compartment 1.3 % of individuals are under treatment, also exposed each of them to antibiotics for 7 days. Antibiotics AbA-AbC-AbF are used in hospital and the community compartments at a proportion (percentage) of 30-40-30; and 75-5-20 respectively. In the basic scenario a single patient treated with only one antibiotic, administered every 8 hours.

### Intensity of the effect of antibiotics on susceptible clinical populations

After each dose administered, all three (bactericidal) antibiotics induce after a decrease of 30% in the susceptible population after the first hour of dose exposure, and 15% in the second hour. These relatively modest bactericidal effects reflect the reduction in antibiotic killing rates of clinical populations when inserted in the colonic microbiota. The antibiotic stochastically penetrates in these percentages of bacterial cells, and those that are susceptible are removed (killed). Therapy is maintained in the treated individual along 7 days.

### Intensity of the effect of antibiotics on colonic microbiota

Antibiotics exert an effect reducing the density of the colonic commensal microbiota, resulting in free-space and nutrients that can benefit the clinical populations. In the basic model, such reduction is 25% for AbA, 20% for AbC, and 10% for AbF.

## Author contributions

FB, CLL, JMS, MC designed the research; MC and FB performed the research; FB, MC, AM, FN, TMC, RC analyzed data; CLL, JMS, MC, RC, RF, FN, and VFL provided computing services, FB and MC wrote the paper.

## Acknowledgements

This work was supported by the European Commission, Seven Framework Program (EVOTAR, FP7-HEALTH-282004) to FB, TMC, VFL, MC; the Instituto de Salud Carlos III of Spain (Plan Estatal de I+D+i 2013-2016, grant PI15-00818; CIBERESP, grant CB06/02/0053) to F.B., the Regional Government of Madrid (InGEMICS-C, S2017/BMD-3691) to TC and FB, SAF2015-65878-R (MINECO, Spain) and PrometeoII/2014/065 (Generalitat Valenciana, Spain) to AM, all co-financed by the European Development Regional Fund, “A Way to Achieve Europe” (ERDF).

---

**Figure SM1.**
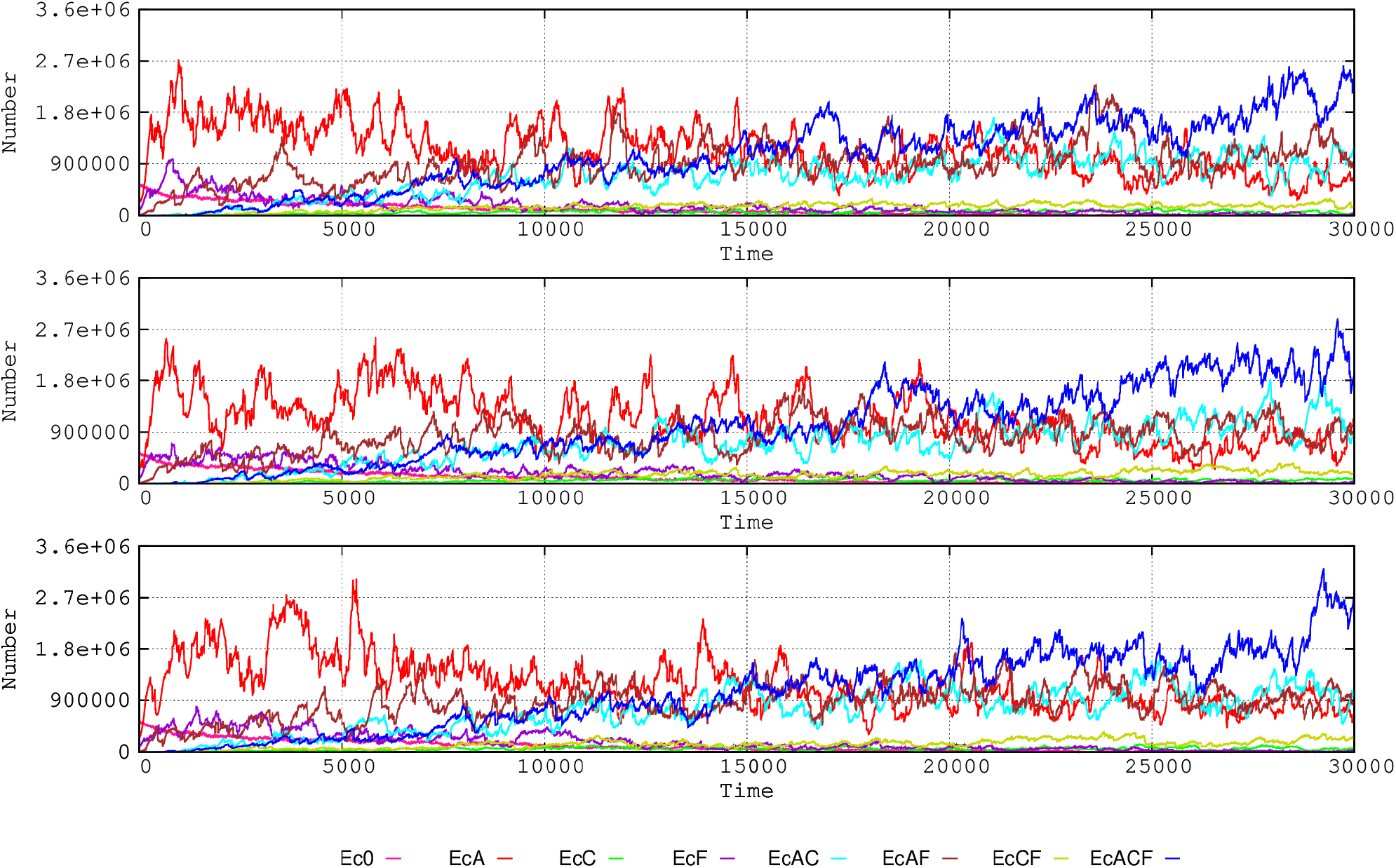
Three consecutive model iterations, in the three panels of the figure, representing the dynamics of *E. coli* resistance phenotypes in the hospital compartment. As the model include several stochastic and probabilistic steps, the results obtained are not entirely identical in replicated runs of the program. However, there are extremely close.

**Figure SM2.**
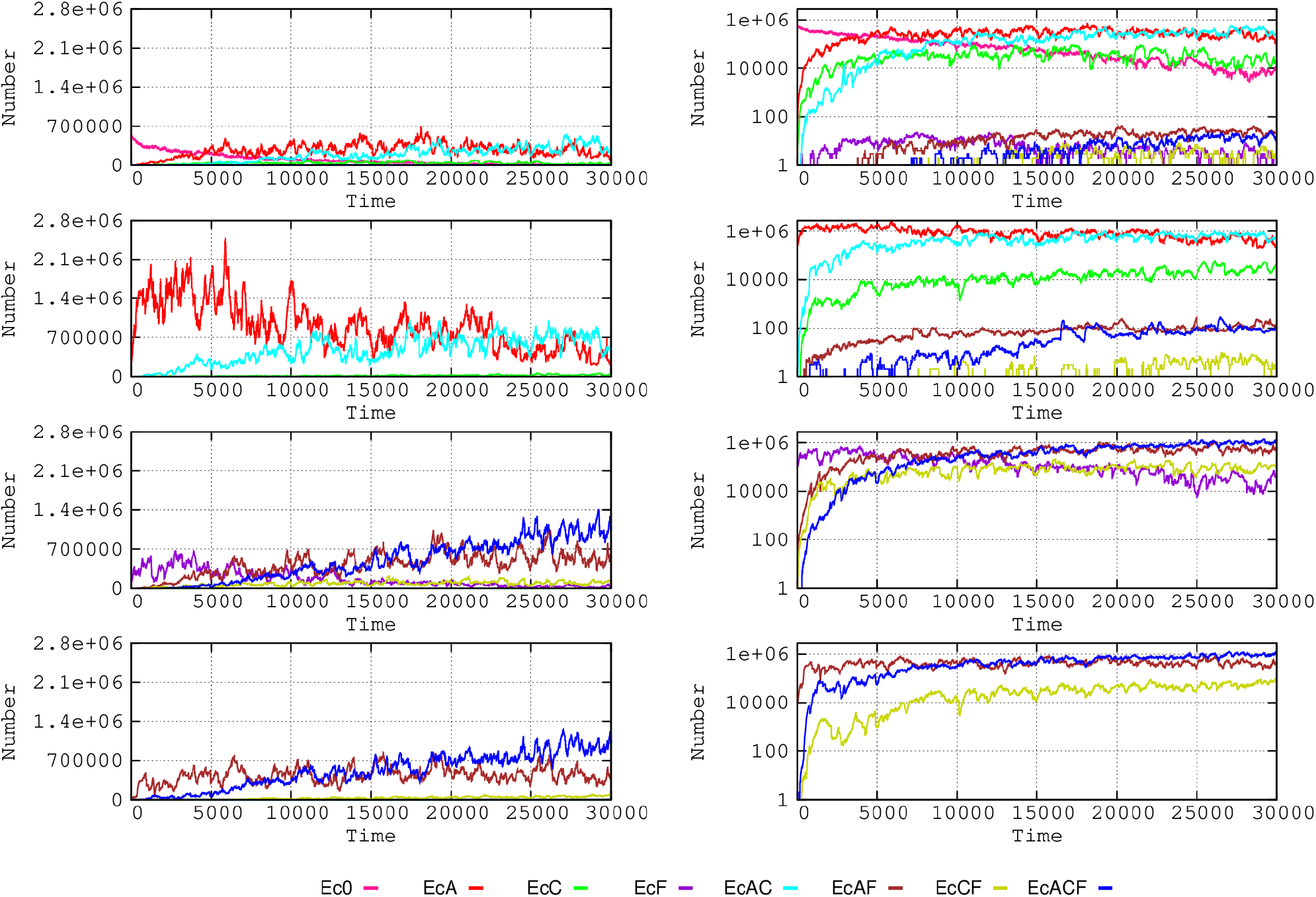
Dynamics of *E. coli* clones starting with different resistance phenotypes in the hospital compartment. On the left half, from top to down, Ecc0 starting without resistance, EccA starting with AbAR; EccF starting with AbFR, and EccAF with AbAR and AbFR. On the right half of the figure, the same in logarithmic representation, allowing to perceive minority phenotypes.

**Figure SM3.**
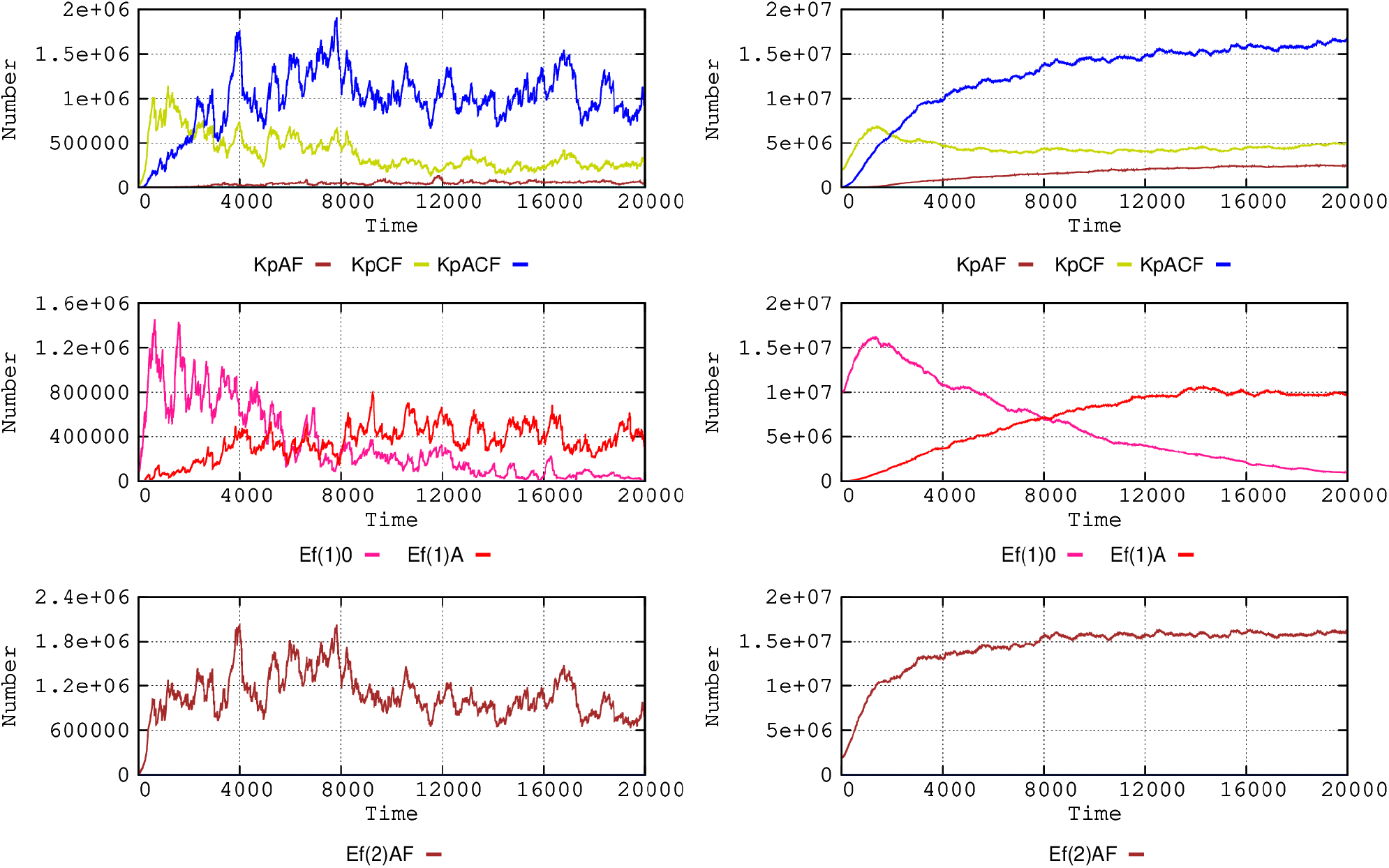
Dynamics of *K. pneumoniae* (top), susceptible *E. faecium* Ef(1) (middle) and Ef(2) AbAR (bottom) in the hospital and community (left and right columns respectively).

**Figure SM4.**
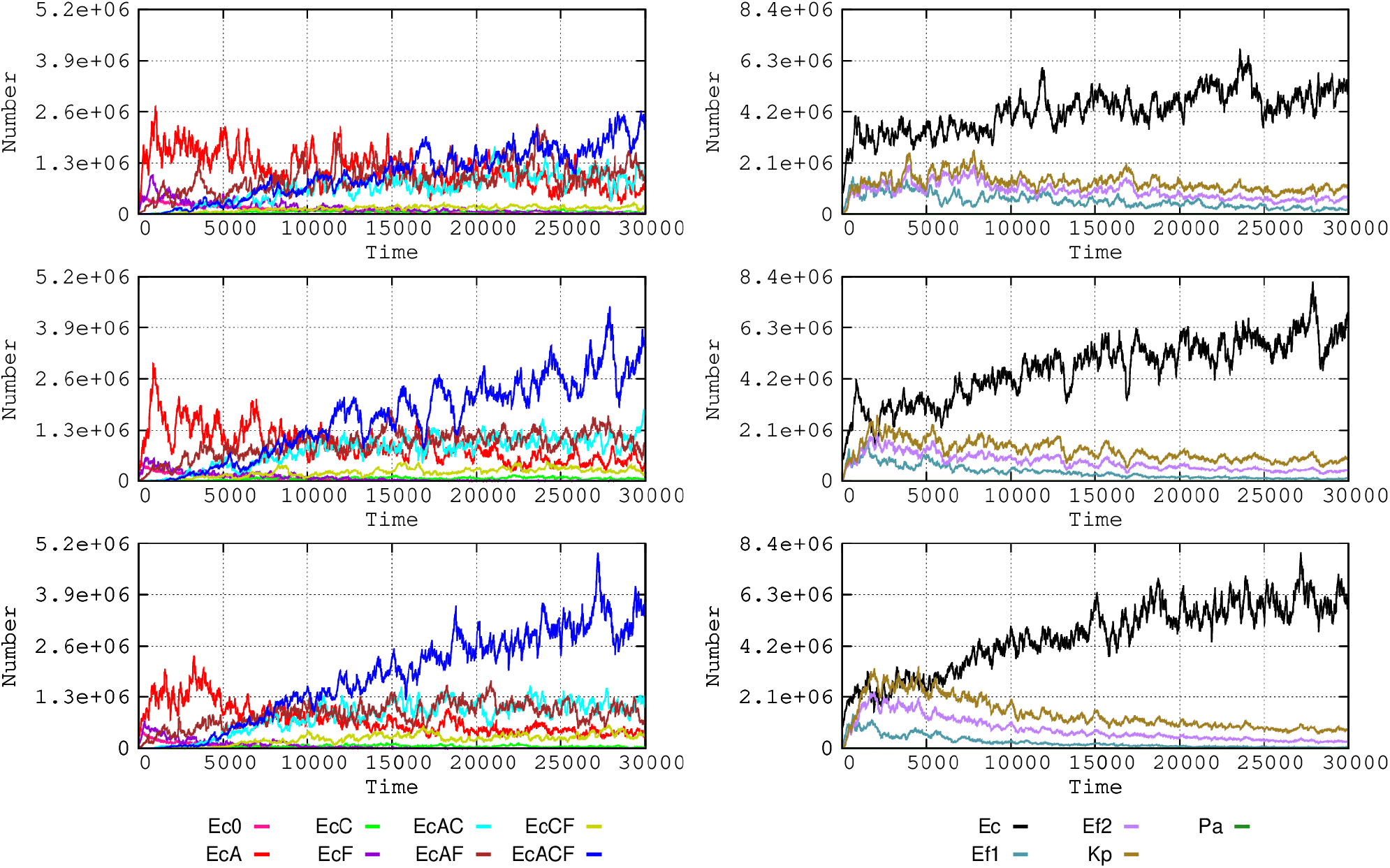
Influence of the size of transmitted bacterial load. On the left half of the figure, *E. coli* phenotypes dynamics in the hospital, when the mean transmitted bacterial load is equivalent to 0.1% (up), 0.5% (mid) or 1% (bottom) of the colonic microbiota. On the right side, evolution of the different species with these transmission loads. Color codes as in Fig 1 and 2.

**Figure SM5.**
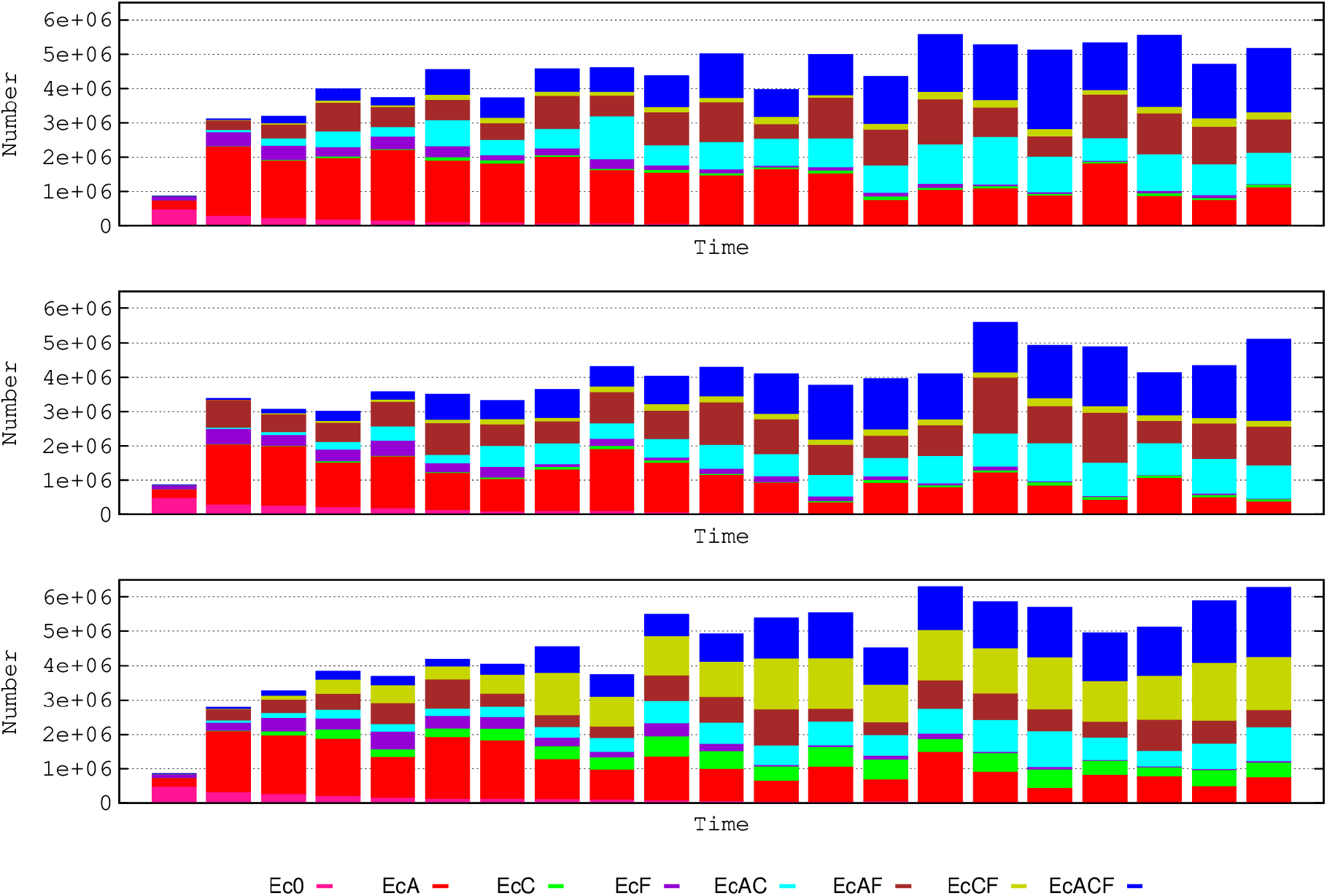
Expected dynamics of hospital-based *E. coli* under the hypothesis that AbCR might provide: 0% of resistance to AbA (upper panel), 10% (mid panel), or 100% (lower panel). Colors as in Fig 1.

